# SigA forms amyloid fibrils that associate with the colicin ColE1 to support the antibacterial activity of *Shigella sonnei* before colonization

**DOI:** 10.64898/2026.07.10.736312

**Authors:** Ahmad Sabbah, Julie Maucotel, Béatrice Roche, Mathieu Erhardt, Lorine Debande, Charlotte E. Chong, Antoine Schramm, Jérémy Fraering, Johana Chicher, Eric Ennifar, Kate S. Baker, Benoit S Marteyn

## Abstract

*Shigella sonnei* is an enteropathogen that causes bacillary dysentery. During the first step of its virulence cycle, it must outcompete the resident microbiota to establish its own niche. Here we report that SigA, the sole SPATE (Serine Protease Autotransporter of Enterobacteriaceae) family member in *S. sonnei*, plays an indirect but central role in this process. A genome-wide analysis showed that the SPATE family includes SigA, Pic, SepA, and Sat. We demonstrated that SigA self-assembles into amyloid fibrils (F-SigA) independently of its protease activity. F-SigA remains associated with the *S. sonnei* surface *in vitro* and *in vivo*. Purified F-SigA fibrils have a diameter of 17.7 ± 3.2 nm, and their amyloid organization was confirmed using specific markers and biochemical methods. F-SigA is secreted into the lumen *in vivo* and localizes to the surface of the colonic epithelium. We found that colicin E1 (ColE1) interacts with F-SigA amyloid fibrils, and that F-SigA–ColE1 complexes display antimicrobial activity that promotes *S. sonnei* competition with other bacteria. Because Pic, another *Shigella* SPATE, also forms amyloid fibrils, we anticipate that this virulence mechanism may be relevant across a wide range of *Shigella* strains and enterobacteria and may serve additional roles during the *Shigella* virulence cycle.

## Introduction

Most, if not all, pathogenic bacteria have deployed colonization strategies relying on their capacity to evolve as bacterial communities producing protective extracellular matrices, which are produced on organ surfaces or within foci of infection. This living mode promotes bacterial genetic diversity, population heterogeneity and protection from the host immune response or other threats such as other bacteria, phages or antibiotics. Extracellular matrices are mainly defined as heterogeneous biofilms containing various cross-linking components such as polysaccharides, nucleic acids, proteins or lipids^1,2^, but may also be structured on single secreted proteins able to form filaments such as type IV pili (*e.g. Neisseria meningitidis*, *Vibrio cholerae*, *Pseudomonas aeruginosa*) ^3^, adhesins ^4^, or amyloid-forming proteins ^5^ .

In this perspective, the colonization strategy of the enterobacteria *Shigella* spp., the causative agents of shigellosis^6,7^, remains unclear, since their ability to produce extracellular matrices has never been reported *in vivo*, although biofilm formation was reported *in vitro*, in the presence of bile salts or endogenous exopolysaccharides ^8–10^ . *Shigella* are pathogenic enterobacteria which colonize and disseminate within the colonic mucosa upon oral challenge; their virulence relies on the expression of several secretion systems (T3SS, T5SS). *Shigella* are amongst the most successful endemic pathogenic enterobacteria and remain a major global health threat due to the rapid emergence of antimicrobial resistant (AMR) strains ^11,12^, and the lack of a commercialized vaccine^13^ .

We hypothesized that the formation of an extracellular matrix may contribute to the dissemination of *Shigella* and that matrix-forming components remain to be identified. *Shigella* are defined as facultative intracellular pathogenic bacteria, although most bacteria evolve extracellularly *in vivo*, within infectious foci formed in the colonic mucosa ^14,15^, then, into the bloodstream, in the context of nutritional deficiency ^16,17^ . *Shigella* species do not produce either flagella or pili, while the *fim* cluster encoding for Type I fimbriae is systematically inactivated in all tested strains ^18^. Until now, the production and secretion of amyloid-forming proteins have not been investigated among *Shigella* strains; curli loci were previously reported to be disrupted in a wide range of strains ^19^ . Various pathogenic bacteria were reported to secrete well-characterized bacterial amyloid fibrils^20^, including *E. coli* or *Salmonella* spp. (curli fibrils, composed of CsgA^21–24^), *Pseudomonas* spp. (FapC, ^25,26^), *Bacillus* spp. (TasA^27^) and *Staphylococcus* spp. (PSMα and PSM β) ^28^ . Those fibrils are formed by monomeric precursors aggregating via β-sheet stacking, resulting in particularly resistant polymeric structures. No homologous protein to these amyloid-forming proteins was identified in *Shigella* spp. On the other hand, EspP and EspC, members of the serine protease autotransporters of enterobacteriaceae (SPATE, T5SS) family were reported to form rope-like fibrils (binding to Congo red and Thioflavin T) in pathogenic *E. coli* species (EHEC and EPEC, respectively) ^29^ . All *Shigella* strains produce and secrete at least one member of the SPATE family, including SigA, SepA and Pic. We hypothesized that these members of the SPATE family secreted by *Shigella* may form amyloid fibrils. We previously demonstrated that SigA (*S. sonnei*) and SepA (*S. flexneri* 5a) are essential for *Shigella* survival to plasma exposure within haemorrhagic colonic mucosa and upon bacteremia, through complement C3 cleavage ^17^ . We have previously reported that SPATEs remain produced and secreted under low-oxygen conditions ^17^, encountered by *Shigella* during its extracellular lifestyle, while *Shigella* T3SS activity is repressed in these conditions ^14,30^. We hypothesized that members of the SPATE family may play a central role in the extracellular lifestyle of *Shigella*; their ability to form fibrils was investigated in this context. In this study, *S. sonnei* and *S. flexneri* 5a were used as models to address these questions.

In the follow-up of previous work on *Shigella* bile salts-induced biofilm formation, we focused our investigation on the early stages of the infection, prior to the invasion of the colon epithelium. In this context, *S. sonnei* competes with the microbiota to establish its own colonization niche. *S. sonnei* produces colicins, including ColE1^31,32^, which plays a central role in this process; the importance of the T6SS was not further confirmed since its identification^33^. The potential interaction between colicins and amyloid fibrils has not yet been reported.

## Results

### *Shigella sonnei* releases extracellular fibrils mainly composed of SigA

We observed by scanning electron microscopy (SEM) that a dense network of fibrils was released *in vitro* by *S. sonnei*, not *S. flexneri* 5a (Fig. 1A); this fibrillar network accumulated in the extracellular compartment and remained associated with the bacterial surface. We hypothesized that these fibrils were amyloid-like complexes, which were previously reported to be released by several other pathogenic bacteria, although not yet described among *Shigella* species. We demonstrated that fibrils released by *S. sonnei* were both stained with the amyloid dyes Congo red (CR) and thioflavin T (ThT), suggesting an amyloid structure (Fig. 1B).

**Figure 1.**
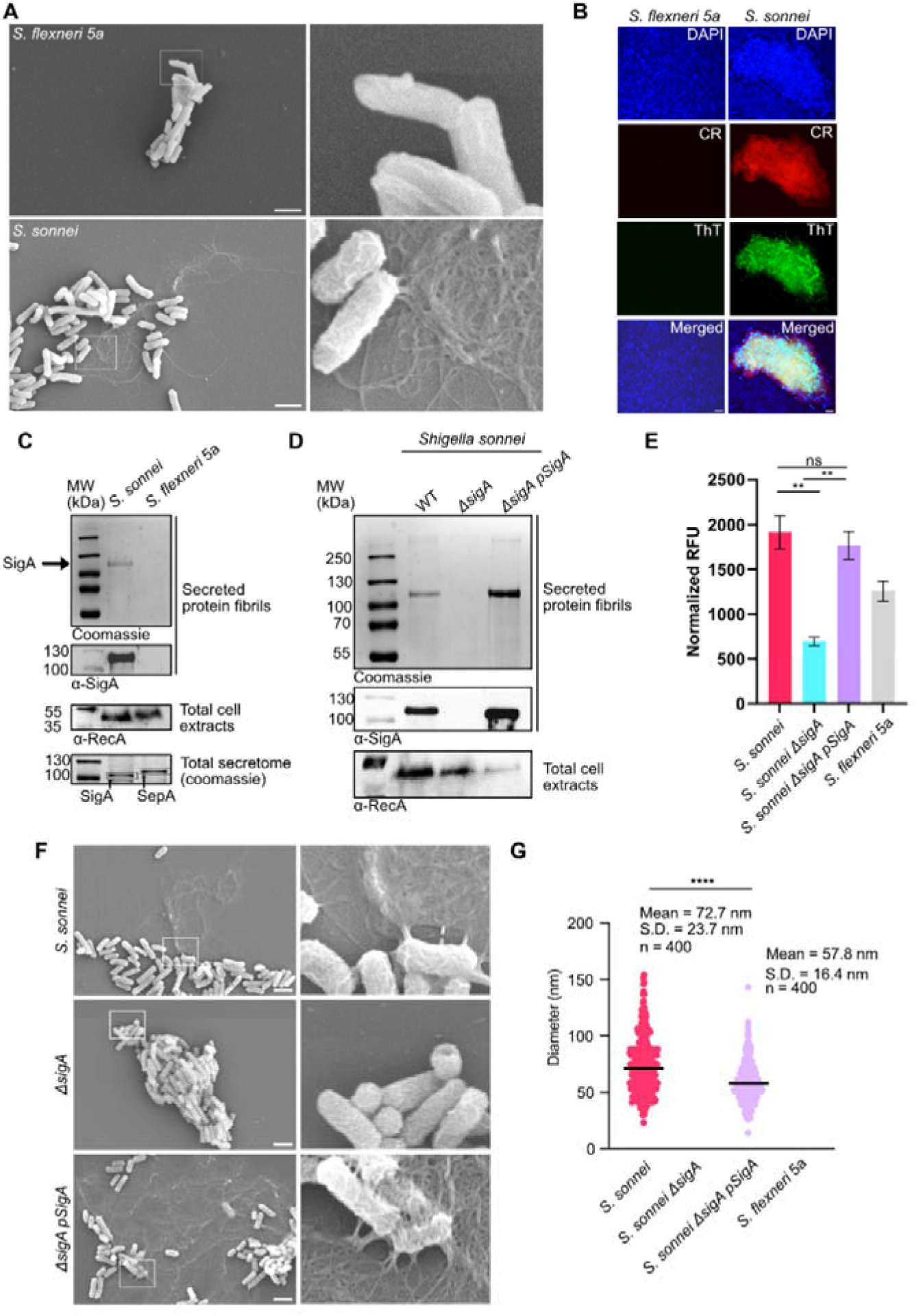
*S. sonnei* SigA oligomerizes into amyloid-like fibrils *in vitro*. (**A**) Extracellular fibrils were detected and analyzed by SEM in *S. sonnei* culture, not *S. flexneri* 5a. Scale bars, 2 μm (**B**) *S. sonnei* extracellular fibrils were labeled with DAPI (blue), congo red (CR, red) and Thioflavin T (ThT, green) and analyzed by epifluorescence microscopy 40X. Scale bars, 10 μm (**C**) Proteins from *S. sonnei* extracellular fibrils were separated on SDS-PAGE together with total cell extracts and total secretome, as controls. Gels were stained with Coomassie, SigA and RecA were detected by western blot. (**D**) As described in (C), secreted protein fibrils were analyzed from *S. sonnei* wild type (WT), Δ*sigA* and Δ*sigA* pSigA strains. (**E**) Quantitative fluorimetric analysis of fibrils secreted by *S. sonnei* wild-type (WT), Δ*sigA* and Δ*sigA* pSigA strains stained with Thioflavin T (ThT) (n=3) ; ns indicates non-significiant, ** indicates p<0.01 (Unpaired Welch’s t-test) (**F**) As described in (B), the release of extracellular fibrils by *S. sonnei* wild type (WT), Δ*sigA* and Δ*sigA* pSigA strains was assessed by SEM. Scale bars, 2 μm. (**G**) Extracellular fibrils diameter (nm) was measured when detected in (A ; F) (n=400). **** indicates p<0.0001 (Mann-Whitney test).

We further aimed to characterize the composition of *S. sonnei* secreted fibrils. To proceed, we applied 1.5M NaCl to separate extracellular fibrils from bacterial cells, as previously described for extracellular matrix extraction ^34^. We further took advantage of amyloid fibril resistance to sarkozyl^35^ and to boiling, to eliminate other sensitive proteinaceous monomers and polymers from the samples, and finally collect fibrils by ultracentrifugation. We confirmed that these fibrils secreted by *S. sonnei* were mainly composed of a major protein, which was identified by mass spectrometry as SigA (Fig. 1C and Supplementary Fig. S1A-B). SigA belongs to the SPATE family, which also includes SepA, Pic and Sat and is the only SPATE expressed by *S. sonnei*. We could confirm by western blot that SigA is detected in secreted fibrils (Fig 1C). Importantly, SigA was also detected as a soluble monomer in the secretome of *S. sonnei*. On the other hand, *S. flexneri* 5a secretes only SepA, which was detected in the total secretome at a comparable level compared to monomeric SigA, but did not assemble into fibrils (Fig. 1C). We confirmed that fibrils were not secreted by a *S. sonnei* Δ*sigA* mutant, as the secretion was restored by complementation (Fig. 1D), confirming that SigA was essential to the formation of these extracellular fibrils. This result was confirmed by quantifying ThT-bound amyloid fibrils in fixed bacteria (Fig 1E), by staining using CR and ThT (Fig. S1C) and by SEM analysis (Fig. 1F). The diameter of the fibrillar components secreted by *S. sonnei* wt was slightly larger compared to those secreted by the complemented strain, although in a comparable range (Fig. 1G, 72.7 ± 23.7 nm vs 57.8 ± 16.4 nm, p<0.0001). The complemented strain, which overexpresses SigA, had a higher density of amyloid fibrils around the bacteria; these results strongly suggested that SigA may assemble into amyloid-like fibrils, hereafter named F-SigA fibrils.

### SigA auto-assembles to form amyloid fibrils

Because SigA fibrils stained positively for both Congo Red and ThT dyes, we next aim at confirming their amyloid nature, using a specific anti-amyloid β antibody (clone WO2), raised against the amyloid cross-β conformation^36^ and previously validated on bacterial amyloid fibrils^37,38^. We confirmed that F-SigA fibrils were detected with WO2 in *S. sonnei* wt and Δ*sigA* pSigA complemented strains, not in Δ*sigA* mutant strain (Fig. 2A), confirming that F-SigA fibrils are amyloid. To identify whether SigA alone was sufficient for amyloid fibrillation, we demonstrated that the overexpression of SigA in *E. coli* HB101 was associated with the secretion of F-SigA fibrils, positively stained with WO2 (Fig. 2B). We hypothesized that the catalytic activity of SigA might play a role in the formation of the fibrils. By overexpressing a catalytically inactive SigA mutant (SigA_S258A_) in *E. coli* HB101, F-SigA fibrils were detected to the same extent (Fig. 2B). This result was further confirmed by SEM analysis (Fig. 2C) and by SDS-PAGE (Coomassie staining and western blot analysis) (Fig. 2D). These results confirmed that SigA is necessary and sufficient for the formation of extracellular amyloid fibrils and that the protease activity of SigA is not essential in this process.

**Figure 2.**
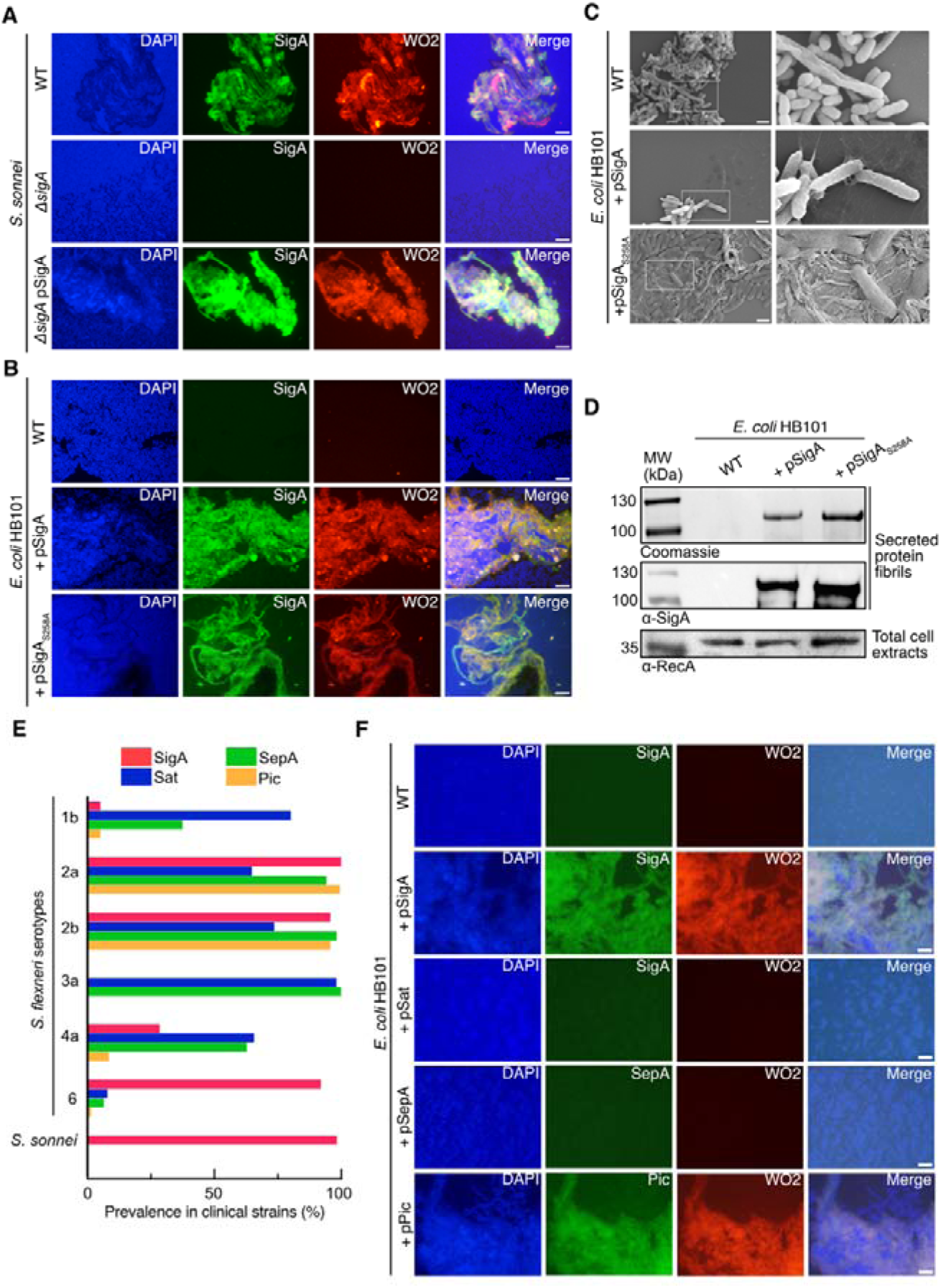
SigA forms amyloid fibrils independently of its catalytic activity. Among other *Shigella* SPATEs, Pic is more prone to form fibrils. (**A**) Secretion of SigA amyloid fibrils by *S. sonnei* wild-type, Δ*sigA* and Δ*sigA* pSigA strains. DNA was stained with DAPI (blue), amyloid fibrils were stained with α-SigA (green) and α-WO2 antibodies (red) and samples were analyzed by epifluorescence microscopy 40X. Scale bars, 10 μm. (**B-C**) Secretion of SigA amyloid fibrils (F-SigA) by *E. coli* HB101 wild-type, pSigA and pSigA_S258A_ strains. (B) Amyloid fibrils were analyzed by epifluorescence microscopy as described in (A). Scale bars, 10 μm. (C) Samples were analyzed by SEM. Scale bars, 2 μm. (**D**) SigA amyloid fibrils (F-SigA and F-SigA_S258A_) secreted by *E. coli* HB101 pSigA and pSigA_S258A_ strains were resuspended in Urea 8M, separated on SDS Page gel and stained with Coomassie or western blotted using a α-SigA antibody. As control, total cell extracts were western blotted using an α-RecA antibody. (**E**) Distribution of SPATEs encoding genes (*sigA*, *sat*, *sepA* and *pic*) in 1,072 genomes of *S. flexneri* (1b, 2a, 2b, 3a, 4a, 6) and *S. sonnei* clinical isolates. (**F**) Detection of amyloid fibrils secreted by *E. coli* HB101 wt, pSigA, pSat, pSepA and pPic strains. DNA was stained with DAPI (blue), SPATEs were stained with specific polyclonal antibodies (green) and amyloid fibrils were stained with the WO2 antibody (red) and samples were analyzed by epifluorescence microscopy 40X. Scale bars, 10 μm.

### Among other *Shigella* SPATEs, Pic is more prone to form amyloid fibrils

Because some *S. flexneri* serotypes secrete SigA, as well as other SPATEs, we aimed to investigate whether other SPATEs would also form amyloid fibrils. It was previously reported that SepA, Pic and SigA were the SPATEs produced by *Shigella* species, although *sat*, the gene encoding for a fourth SPATE, Sat, was identified in a significant number of clinical strains ^39^. We analysed the distribution of the SPATE genes (*sigA*, *sepA*, *pic* and *sat*) in 1,072 *S. sonnei* and *S. flexneri* genomes from the Global Enteric Multicentre Study (GEMS) (Supplementary Table S4). GEMS was a large, multi-country case-control study investigating the causes of moderate to severe diarrhoea in children under five years old in low- and middle-income countries ^40^. We confirmed that all *Shigella* species contained at least one SPATE encoding gene in their genome. *sigA* was the lone member of the family detected in *S. sonnei*, as expected ^17^ and was the most prevalent in *S. flexneri* 6 isolates. *sepA* and *sat* were detected in most *S. flexneri* species (1b, 2a, 2b, 3a, and 4a) and in a minority of *S. flexneri* 6 isolates. *pic* was mainly restricted to *S. flexneri* 2a and 2b (Fig. 2E). As the structure of SPATEs is well-conserved, we evaluated their self-assembly ability to form fibrils. Using the heterologous ovexpression strategy in *E. coli*, combined with WO2 staining, we demonstrated that Pic was more prone to form amyloid fibrils *in vitro*, as compared to SepA or Sat, and to the same extent as SigA (Fig. 2F). This result was confirmed on SDS Page (Supplementary Fig S2). This result was unexpected considering the structural homology of SPATEs ^17^, including Sat (Fig. S3), and suggests that specific amino acid sequences conserved in SigA and Pic are essential for their auto-assembly into amyloid fibrils.

We further aimed at optimizing the purification of F-SigA fibrils and defining their biochemical organisation.

### Purification and biochemical characterization of F-SigA amyloid fibrils

To proceed with the biochemical analysis of F-SigA fibrils, we used a dual approach, combining an *in-silico* structure prediction strategy with an *in vitro* biochemical analysis, which required the production of F-SigA at a high level of purity and at high yield *in vitro*. As a preliminary approach, we first explored the theoretical SigA potential for oligomerization at the molecular level using *in silico* prediction methods. First, using AlphaFold-3, we address the question of whether a rigid-body SigA multimerization model could be compatible. Based on the prediction of SigA dimer interfaces and their subsequent combination (Supplementary Fig. S3A), six putative polymeric structures free of inter-molecular steric clashes were predicted (Supplementary Fig. S3B). This initial approach tended to confirm the capacity of SigA to auto-assemble, although the structure prediction tools used for this initial approach are limited to rigid-body models. Thus, given SigA as the core component of *S. sonnei* released amyloid fibrils (Fig. 2), we further investigated structural features within the passenger domain of SigA, which may be involved in amyloid-like structural rearrangements. To proceed, SigA sequence analysis with AmylPred2 revealed multiple regions of high amyloidogenic propensity. Because SigA and Sat are structurally very similar (Fig S4A), their opposing behaviour in terms of amyloid fibrillation sparked our interest. AmylPred2 programs predicted numerous amyloid hotspots of interest on SigA (Fig S4B) but also Sat (Fig S4C) with a consensus well above the threshold. Thus, we hypothesized that SigA amyloidogenic regions might be more spontaneously exposed than Sat to promote amyloid fibril formation.

To go beyond these predictive results, we purified and characterized F-SigA *in vitro* using classical biochemical methods. The production of pure F-SigA was technically challenging due to its tight binding and association with bacteria in culture. A specific method was optimized to proceed, consisting in the induction of the fibrillation (18h at 37°C) of pre-secreted monomeric SigA in a culture medium devoid of bacteria, following a series of filtration steps, sonication and concentration by ultracentrifugation (Fig. 3A, and see Methods). The high degree of purity of F-SigA obtained through this procedure was confirmed on SDS-PAGE (Fig. 3B). The structure of these ultrapure F-SigA fibrils was analyzed by TEM (Fig. 3C); their diameter was quantified (17.7 ± 3.2 nm) and appeared to be significantly lower than the large bundles of fibrils associated to bacteria (Fig. 1G). To characterize F-SigA we first conduct a Congo Red assay, which displayed a shift in the peak of the F-SigA spectrum compared to monomeric SigA or Congo Red alone (Fig. 3E). This result was consistent with an increased Thioflavin T binding to F-SigA compared to monomeric SigA (Fig. 3F). Furthermore, as other amyloid fibrils, we confirmed that pure F-SigA amyloid fibrils were stained with Congo Red, Thioflavin T and α-WO2 specific antibody, and do not contain DNA, as reported by a negative staining with DAPI (Fig. 3G).

**Figure 3.**
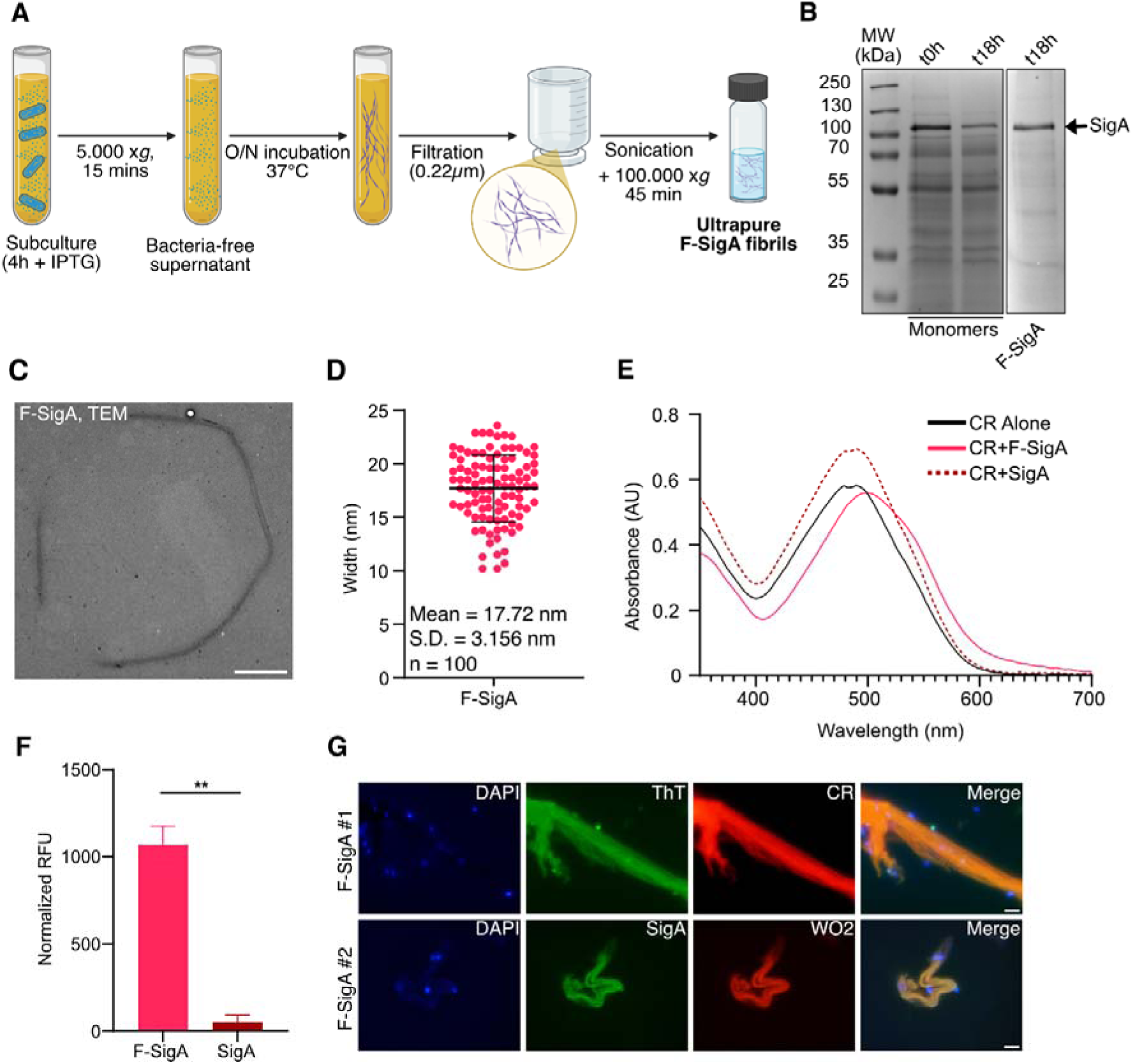
Biochemical characterization of ultrapure F-SigA. (**A**) Experimental procedure allowing the formation and purification of pure SigA amyloid fibrils. Created in BioRender (https://BioRender.com/dcb86fk)(B) Purified F-SigA were resuspended in Urea 8M, while monomers at t0 and t18h were concentrated 350x and separated on SDS-PAGE gel and stained with Coomassie. (**C**) Purified F-SigA were analyzed by TEM (**D**) Purified F-SigA width was measured on different fibrils (n=100). Result is expressed as Mean and S.D(**E**) Representative spectrophotometric analysis of SigA and F-SigA stained with congo red (CR). (**F**) Quantitative fluorimetric analysis of SigA and F-SigA stained with Thioflavin T. Results are expressed as mean ± S.D. (n=3). ns indicates non-significant, ** indicates p<0.01 (Unpaired Welch’s t-test) (**G**) Purified F-SigA (samples #1 and #2) stained with DAPI (blue), ThT or SigA (green), CR or WO2 anti-amyloid beta (red) by epifluorescence microscopy using a x100 objective. Scale bar = 2 µm.

In conclusion, we developed a dedicated method to purify F-SigA, and we confirmed using complementary approaches that monomeric SigA auto-assembles to form amyloid fibrils *in vitro*. We further aimed at defining the contribution of F-SigA and associated amyloid matrix to the virulence of *S. sonnei*, and to clarify its interrelationships with bile salts-induced biofilm.

### Contribution of F-SigA to the extracellular lifestyle of *S. sonnei*

We first demonstrated that F-SigA was released by *S. sonnei in vivo*, using the ascorbate-deficient guinea pig model of shigellosis ^16^. We could detect amyloid matrices co-stained with α-SigA and α-WO2 antibodies in the vicinity of *S. sonnei* wild-type strains, within the colonic lumen, close to the epithelium surface. This result was confirmed using a *S. sonnei* Δ*sigA* pSigA strain, which overexpresses SigA (Fig. 4A). No F-SigA was detected upon infection with the *S. sonnei* Δ*sigA*,, which was previously shown to be defective for colonization ^17^.

**Figure 4.**
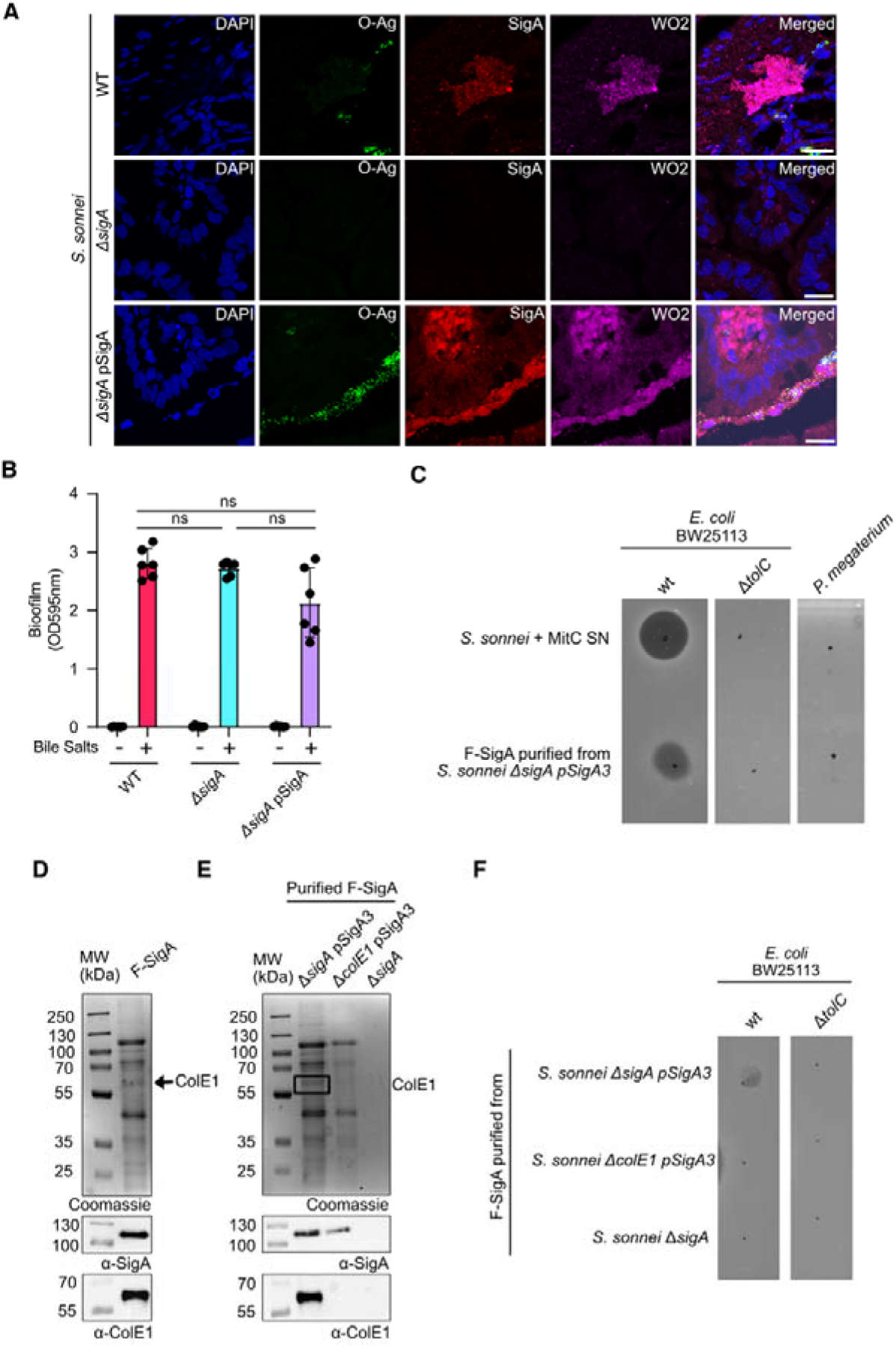
F-SigA amyloid fibrils are secreted by *S. sonnei in vivo* and possess bactericidal activity by association with ColE1. (**A**) Guinea pig colons infected by *S. sonnei* WT, Δ*sigA*, and Δ*sigA* pSigA3 strains. 48h p.i. samples were collected and stained with an α-*S. sonnei* O-Ag (O-Antigen) antibody (green), an α-SigA antibody (red), an α-WO2 antibody (magenta), DNA was stained with DAPI (blue). Bars, 50 µm. Representative results from n > 3 animals per condition. (**B**) Bile salt-induced biofilms formed by *S. sonnei* WT, Δ*sigA*, and Δ*sigA* pSigA strains (24 h in LB+0.25% glucose supplemented or not with 0.4% bile salts) (n=3). ns indicates p>0.05 (Paired Welch’s t-test). (**C**) *E. coli* (Gram-negative) and *P. megaterium* (Gram-positive) lawn killing assays on LB soft agar with culture supernatant (SN) of *S. sonnei* strain treated with mitomycin C (MitC, 0.5 µg/mL) or F-SigA purified from *S. sonnei* Δ*sigA* pSigA strain. (**D**) F-SigA purified from *S. sonnei* Δ*sigA* pSigA3 were separated on SDS-PAGE gel and stained with Coomassie or western blotted using α-SigA and α-ColE1 antibodies. (**E**) F-SigA purified from *S. sonnei* Δ*sigA* pSigA3, Δ*colE1* pSigA3 or Δ*sigA* were analyzed as described in (D). (**F**) *E. coli* BW25113 WT and Δ*tolC* lawn killing assays on soft agar against F-SigA from strains described in (E). Our results provide new insight into the function of SPATEs during the initial stage of the infectious process.

Amyloid fibrils are frequently associated with bacterial biofilm scaffolding. *Shigella* biofilm formation remains discussed; only bile salts were shown to induce biofilm formation in *S. sonnei in vitro*^8^. As the expression of *sigA* was reported to be induced by bile salt-associated regulators^41^, we hypothesized that their formation may be linked to the secretion of F-SigA. We confirmed that *S. sonnei* wild-type strain formed a biofilm, only in the presence of bile salts, and we demonstrated that the level of production by *S. sonnei* Δ*sigA* and complemented strains was comparable (Fig 4B). This result confirmed that F-SigA was not essential to *S. sonnei* bile salt-dependent biofilm formation.

As we detected F-SigA within the colonic lumen (Fig. 4A), we hypothesized that it may be important for the extracellular lifestyle of *S. sonnei* in this microenvironment and may contribute to the competition of *S. sonnei* with the resident microbiota. Indeed, we confirmed that F-SigA purified from *S. sonnei* Δ*sigA* pSigA3 (IPTG-inducible *sigA*) has a bactericidal activity on *E. coli*, used as Gram-negative bacteria model, but not on *Priestia megaterium*, used as Gram-positive bacteria (Fig. 4C). We further demonstrated that F-SigA bactericidal activity was abolished in an *E. coli* Δ*tolC* mutant strain, suggesting that a colicin mediated this activity ^42^. *S. sonnei* CIP106347 was previously shown to secrete a single colicin, ColE1^43^. This hypothesis was supported by the similar results obtained with culture supernatants of a *S. sonnei* strain treated with mitomycin C (inducing ColE1 secretion) (Fig. 4C), or *S. sonnei* Δ*colE1* mutant and complemented strains treated similarly, or purified ColE1 (Supplementary Fig. S5). The association between F-SigA and ColE1, suggested by these results, was confirmed by Coomassie staining and western blot analysis on F-SigA purified from the *S. sonnei* Δ*sigA* pSigA3 (Fig 4D). We further confirmed that ColE1 was absent from F-SigA purified from a *S. sonnei* Δ*colE1 and* Δ*sigA* mutants, as controls (Fig. 4E). We further demonstrated that F-SigA purified from *S. sonnei* Δ*colE1* mutant did not affect *E. coli* growth, confirming the essential role of ColE1, through its association, on F-SigA bactericidal activity.

## DISCUSSION

The production of extracellular matrices or biofilms by *Shigella* has long been debated. A bile-salt-induced biofilm formation was previously reported *in vitro* ^8^, although it has been challenging to confirm this result *in vivo*, due to the lack of optimal animal models of shigellosis. We have demonstrated here that at least two members of the SPATE family, SigA and Pic, have the capacity to auto-assemble post-secretion onto amyloid fibrils *in vitro*. These results are consistent with a previous report highlighting the ability of two SPATEs (EspC and EspP) secreted by pathogenic *E. coli* strains (EPEC and EHEC, respectively) to form rope-like structures that stain positively for the amyloid dyes Congo Red and Thioflavin T ^29^. F-SigA are, to our knowledge, the first secreted amyloid fibrils identified in *Shigella* strains. Among other amyloid proteins, several genes belonging to the curli operon were identified in the genome *S. flexneri* 2A (2457T)^44^, although it was reported that the curli locus was lost by *Shigella* strains^19^. Although SigA and, by extension, F-SigA, were not essential for biofilms when exposed to bile salts, further investigation will be required to define whether F-SigA formed by *S. sonnei in vitro* and *in vivo* may be integrated in a biofilm. The question of a biofilm could be extended to the association of F-SigA with other components, such as exopolysaccharides, lipids or nucleic acids, scaffolding a heterogeneous bacterial population. More studies on biofilm formation beyond bile salts are required to address this question. And beyond, it will be of interest to evaluate to what extent amyloid fibrils released by *Shigella* may illustrate the concept of matrixome, beyond the classical definition of biofilms^2^ .

In this princeps study, the F-SigA matrix was observed within the colonic lumen, at the epithelium surface; further investigation will be required to define its formation during subsequent steps of the infectious process, first, within the colonic mucosa during the formation of foci of infection, and beyond, in the blood circulation, during bacteremia, as we previously demonstrated that SigA is essential for *S. sonnei* survival to plasma exposure^17^. The outcome of the interaction between F-SigA amyloid fibrils and immune cells will be particularly interesting to evaluate their impact on their antimicrobial and immune functions. Further, the interaction of F-SigA with extracellular DNA is of great interest and will have to be further defined, as amyloid-DNA complexes have been shown to exhibit specific and important immunomodulatory functions, particularly in the context of curli secretions ^45,46^.

Conversely, if all *Shigella* species express at least one SPATE, we reported from a global genomic study that several strains are devoid of *sigA* and *pic* (e.g. *S. flexneri* 1b or 3a, Figure 2), and may, according to our results, be defective for the production of amyloid fibrils. Further investigation will be required to evaluate if other SPATEs may form amyloid fibrils in other environmental or pathophysiological conditions or if other alternative systems may be expressed as a compensation to promote the formation of extracellular matrices and *Shigella* adhesion. Many genes encoding potential matrix-forming virulence factors located in the virulence plasmid are inactivated, potentially by pathoadaptive mutations^47^ . As a consequence, not all *Shigella* strains express adhesins on their surface, although an adherence domain was identified in the IcsA passenger domain^48,49^. A multivalent adhesion molecule (MAM) was identified in *S. sonnei,* but was not further reported in other *Shigella* species ^50^.

Unexpectedly, we demonstrated that the protease activity of SigA was not essential for its ability to auto-assemble. Conversely, the impact of the fibrillation on the modulation of SigA protease activity will have to be clarified, as those fibrils may be formed in a concentration-based manner or to temporarily inactivate SigA in a structurally resistant state. Virulence factor regulation through amyloid fibrillation was shown in Microcin E492 of *Klebsiella pneumoniae,* whereas that bacteriocin was shown to be stored as a toxic reservoir^51^. Furthermore, toxin self-regulation was shown in the context of the cytotoxin listerolysin O produced by *Listeria monocytogenes*, where amyloid aggregation inhibited the toxic activity of the protein^52^.

We report here a first contribution of SigA amyloid fibrils to *Shigella* colonization strategy through their interaction with the colicin ColE1. The association of ColE1 with F-SigA into amyloid fibrils released by *S. sonnei* raises additional questions regarding the mode of ColE1 secretion in these physiological conditions. In addition, as we demonstrated that ColE1 was active on E. coli while associated with F-SigA, the affinity of this interaction and its regulation will have to be further investigated to define how ColE1 reaches TolC, its receptor in targeted bacteria, in this context.

Colicins were previously shown to be conserved among epidemiologically successful *S. sonnei* clinical strains^31,32^. Several bacteriocins have been previously reported to form amyloids, such as Blon_0434 from *Bifidobacterium longum* ^53^ or microcins from *K. pneumoniae*^54^ or from *Acinetobacter baumannii* ^55^. However, the interaction of bacteriocin, including colicins, with an amyloid fibril has not yet been described. The biochemical aspects of this interaction will have to be determined to better appreciate its stability during the infectious process. The incorporation of this potent antimicrobial protein secreted by *S. sonnei* within the amyloid matrix is anticipated to promote the colonization of the colonic lumen. It was previously reported that within *E. coli* biofilms, the production of colicins was specifically increased (as reported for colicin R ^56^ and colicin E2 ^57^). Whether the production of colicins by *S. sonnei* is modulated in the context of F-SigA amyloid fibrils will have to be characterized.

In conclusion, the ability of SPATEs to auto-assemble into amyloid fibrils sheds new light on the contribution of this protein family to *Shigella* and other enterobacteria virulence. SPATEs have been previously shown to play multiple roles, associated with the cleavage of multiple targets, during *Shigella* virulence cycle; from mucus degradation, to the destabilization of the epithelium, to the subversion of immune cell functions, to the inhibition of complement activation (as reviewed in ^58^). The respective contribution of the amyloid forms of SPATEs to these different functions will have to be deciphered *in vitro* and *in vivo*.

## MATERIALS AND METHODS

### Bacterial strains and plasmids

All strains and plasmids used in this study are listed in Table S1. *S. sonnei* WT strain was obtained from the Pasteur Institute Collection (CIP106347). The *S. sonnei* Δ*sigA* strain belongs to the laboratory collection ^17^ . *S. flexneri* 2a 2457T, *S. flexneri* 5a M90T-Sm, *E. coli* HB101 and *E. coli* DH5α wild-type strains belong to the laboratory collection. Uropathogenic *E. coli* CFT073 was provided by Eric Oswald’s lab. *Priestia megaterium* (formerly *Bacillus megaterium*) was kindly given by Isabelle Caldelari’s lab.

*S. sonnei* Δ*colE1imlys* mutant strain was constructed with the lambda-red recombination method ^59^. In brief, *colE1, immunity* and *lysis* genes on the pColE1 plasmid were replaced by a chloramphenicol resistance cassette, which was amplified from the pKD3 plasmid using primers listed in Table S3.

Plasmid transformations in *Shigella* were performed by electroporation and, in *E. coli* were performed using TSS-competent *E. coli.* SPATE- and ColE1-expressing plasmids were constructed using the HiFi DNA Assembly (NEB) technique. pBBR1-MCS4 was used and modified for the expression of SPATE encoding genes (the plasmid was linearized by PCR, excluding the Lac promoter, the MCS4 multiple cloning site and the *lacZ*α gene). SPATE encoding genes were amplified from lysates of wild type strains of *Shigella sonnei* (*sigA*), *S. flexneri* 2a (*pic*), *S. flexneri* 5a (*sepA*) and Uropathogenic *E. coli* (*sat*) using 60bp primers (25 complementary bp + 35 bp overhang homologous to amplified pBBR1’s ends). Once constructed, the pBBR1-*sigA* (pSigA) plasmid served as the backbone for the cloning of *sepA*, *pic* and *sat* (after removing *sigA* coding sequence, while retaining *sigA* native promoter and terminator). Of note, pBBR1-pic (pPic) was constructed with a modified version of the *pic* gene, lacking set toxin-encoding sequence, using alternative *pic* codons in the *set* region. The plasmid pUC19-*sigA* was kindly provided by Dr. Mario Meza Segura ^60^. For pBR03 (pET28-ColE1-His6), the pET28a plasmid was used as a backbone and was linearized by PCR. *colE1* was amplified from pColE1_CIP106347_ of wild type *Shigella sonnei* strain using 60bp primers (25 complementary bp + 35 bp overhang homologous to amplified pET28a’s ends). For pColE1 (pCU19_P*colE1*_*colE1immlys*), pUC19 was used as a backbone and linearized by PCR, while the *colE1* promoter and *colE1, immunity* and *lysis* genes were amplified from pColE1_CIP106347_. PCR products were treated with DpnI FastDigest (Thermo Fisher Scientific) for 1 hour at 37°C then purified using PCR Clean Up columns (MN). 0.05 mol linearized vector was mixed with 0.1 mol inserts, HiFi Assembly Master Mix 2x and milliQ water. Assembly was performed at 50°C for 20 minutes, and 2 µL were transformed in TSS-competent^61^ DH5α cells. Plasmid pBBR1-SigA_S258A_ was generated using site-directed mutagenesis combined with HiFi DNA Assembly (NEB), using primers listed in Table S3. Briefly, the parental plasmid was linearised by PCR in 2 fragments with homologous ends using a pair of mutagenic primers and a pair of fully homologous primers located on the opposite side of the plasmid. PCR products were treated with DpnI Fast Digest for 1 hour to remove parental plasmid, and purified using PCR Clean Up. The 2 fragments were assembled using HiFi assembly and transformed as described previously.

All PCRs were performed using SuperFi II polymerase (Thermo Fischer Scientific). All transformants were screened by PCR on colony using Taq polymerase. Validated plasmids were purified using MiniPrep Kit (MN) and verified by Whole Plasmid Sequencing (Eurofins).

### Microscopy analysis of purified or bacterial amyloid fibrils

Bacterial cultures were grown O/N at 160RPM 37°C in LB-Miller (1% Tryptone, 1% NaCl and 0.5% Yeast Extract) medium. Cultures were centrifuged at 500g for 5 minutes to pellet mostly aggregated bacteria or structure-associated bacteria. The pellet was fixed in 4% PFA in cold PBS for 30 minutes, followed by washing with cold PBS.

For immunofluorescence, samples were washed with cold PBS mixed with 0.1% Triton and 1% BSA, then incubated in the same blocking solution supplemented accordingly with 150 µM Congo Red(Sigma), 150 µM Thioflavin T (Sigma), rabbit anti-SigA ^17^ conjugated with Alexa Fluor 568 lightning link kit (abcam), mouse anti-amyloid beta clone WO2 (Thermo Fischer) conjugated with Alexa Fluor 647 lightning link kit (abcam), mouse anti-*S.sonnei* OAg (described in this paper) conjugated with ChembrightPlus Coralite Plus 488 conjugation kit (ProteinTech). Samples were imaged with an Axioskop two light microscope (Zeiss, Germany) equipped with an Optikam Pro6 digital Camera (Optika, Italy), using a 40x (Zeiss EC Plan-Neofluar 40x/0.75) or 100x objective (Plan-NEOFLUAR 100X 1.3NA Ph3 oil).For Scanning Electronic Microscopy (SEM), samples were fixed in 3.7% paraformaldehyde for 30 mins at room temperature. After washing, samples were post-fixed for 2 h with 1% osmium tetroxide and dehydrated through a graded ethanol series (50%, 70%, 90% and 100%) for 30 min at each step. Samples were then deposited onto glass coverslips and treated with hexamethyldisilazane. After solvent evaporation, dried samples were mounted on SEM stubs using carbon adhesive tabs and sputter-coated with gold. Observations were performed using a ZEISS Sigma 300 HV scanning electron microscope operated at 10 kV with a secondary electron detector.

For Transmission Electronic Microscopy (TEM), a drop of the sample preparation was deposited onto a carbon film–coated 300-mesh nickel grid held with fine forceps. After allowing particles to adsorb to the carbon film for 5 min, excess liquid was removed using filter paper. The grid was then negatively stained with 0.2% uranyl acetate. After a few seconds, excess stain was blotted off with filter paper. Observations were carried out using a ZEISS Sigma 300 HV scanning electron microscope operated at 30 kV in a scanning transmission electron microscopy (STEM) mode.

Imaged purified fibrils were processed similarly to bacteria for imaging, but without fixing with PFA, and centrifugations were performed at 100,000 x*g* for 20 minutes. Quantification of fibril width in all conditions was performed using ImageJ.

### Isolation of fibrils from bacterial cultures

Bacterial cultures were grown O/N at 37°C in LB medium. Cultures were centrifuged to pellet the bacteria at 3200g 4°C for 15 minutes. The supernatant containing secreted proteins was filtered through 0.22 µm nitrocellulose filter, and proteins were precipitated with 35% ammonium sulphate at 4°C for 30 minutes. Precipitated proteins were collected by centrifugation at 7500x *g* 4°C for 30 minutes.

The bacterial pellet was resuspended in 1.5M NaCl by pipetting and vortexed for 30 seconds. Following 3200 x *g* centrifugation for 15 minutes, the fibril-containing supernatant was collected in a clean tube while the bacterial pellet was washed in PBS, then cell lysates were prepared at 95°C for 15 minutes. The fibril-containing supernatant was supplemented with 1% sarkosyl (Sigma) and briefly vortexed before incubation at 95°C for 15 minutes. The suspension was centrifuged at 5000g for 5 minutes to remove initial cell debris and the supernatant was further ultracentrifuged at 100,000 x *g* 4°C for 45 minutes. The sarkosyl-insoluble pellet was resuspended in 8M Urea later supplemented with 1:6 Laemmli 6X and boiled at 95°C for 10 mins. Bacterial cells were boiled for 95°C for 10 minutes, centrifuged at 17000g for 10 minutes to remove cell debris, and the supernatant was collected and mixed with 1:6 Laemmli 6X. All fractions were analysed on 10% SDS-PAGE gels stained with Instant Blue Coomassie (Euromedex) or transferred to nitrocellulose membranes (Amersham) using the transblot standard program (BioRad) and stained with 1 µg/mL rabbit anti-SigA ^62^, rabbit anti-RecA (ProteinTech), and detected with HRP-conjugated goat anti-rabbit (ProteinTech).

### Purification of amyloid fibrils for functional biological assays

Overnight cultures of a strain expressing pSigA3 (pUC19-*sigA* inducible with IPTG) were subcultured from OD600=0.05 to OD=2 in LB supplemented with Ampicillin 100 µg/mL and IPTG 1 mM. Bacteria were centrifuged at 6000g 4°C for 15 minutes. The supernatant was filtered through 0.22 µm filtration units onto clean glassware. The filtered supernatant was supplemented with Kanamycin 50 µg/mL (to inhibit *S. sonnei-*derived strains) or Streptomycin 50 µg/mL (to inhibit *E. coli* HB101 strains) to avoid any contamination by the producing strain and incubated at 37°C 160 RPM for 18 hours to allow fibrillation.

Amyloid fibrils were collected by re-filtering the incubated bacteria-free medium. The fibrils captured by the 0.22 µm membrane were washed by filtering 2 volumes of NaCl 1.5M and 1volume of deionized water. The fibrils were collected in 10 mL PBS, sonicated at 120V for 5 seconds followed by 5 secs resting on ice (5 cycles). The bigger fibrils were collected by centrifugation at 1000g and 20,000 x g for 20 mins, while the smaller fibrils were collected at 50,000 x g and 100,000 x g for 20 mins each. Detergent-free and boiling-free amyloid fibrils collected at lower centrifugation speed were used for general purpose, while fibrils fractionated at higher centrifugation speed were used for microscopy-related experiments.

To visualise the dynamics of fibrillation, 50 mL were collected at t0, corresponding to the filtered supernatant before the overnight incubation, and 50 mL were collected 18 hours later. Supernatants were concentrated to 150 µL using centricon ultracentrifugation 50-kDa cut-off tubes, followed by boiling with Laemmli 6x at 95°C for 10 minutes. Fibrils were resuspended in urea 8M and boiled for 10 minutes, before adding Laemmli 6X and boiling further for 5 minutes. Samples were migrated on 10% SDS-PAGE and either stained with Instant Blue Coomassie (Euromedex) or analysed by western blot as described before, using 1 µg/mL rabbit anti-SigA ^62^, or rabbit anti-ColE1 (this study) antibodies.

### Congo Red and fluorimetric analysis of amyloid fibrils

Bacterial cultures were collected and fixed exactly as with microscopy. Fixed bacteria OD600 was measured then they were resuspended in PBS supplemented with 20 µM ThT and incubated at room temperature in the dark for 30 minutes. Fluorimetric analysis was performed with DeNovix ds-11 fx+ using 350 nm excitation. Raw RFU was normalized by subtracting the values of the control ThT and adjusting with the OD600 values.

Bacteria-free fibrils, or purified SigA monomers, were quantified by nanodrop at OD280 to quantify proteins. They were then mixed with 20 µM Congo Red or 20 µM ThT and incubated at room temperature for 30 minutes. Congo Red spectrum was performed using DeNovix ds-11 fx+ “UV-Vis” program. Samples without Congo Red were also measured to normalize. ThT fluorimetric analysis was performed with ds-11 fx+ using 350 nm excitation. Raw RFU was normalized by subtracting the values of the control ThT and adjusting with the OD280 values

### Production of α-ColE1 and α- *S. sonnei* O-Antigen antibodies

#### Anti-ColE1

*E. coli* Nico21 strain was transformed with pET28-ColE1-His6 plasmid. Overnight culture was inoculated into LB medium supplemented with Kanamycin 50 µg/mL and grown at 37□°C until OD600 reached 0.6. Then, protein production was induced by the addition of 0.5 mM IPTG for 4□h at 37□°C. Cell pellet was re-suspended in lysis buffer (50 mM Tris-HCl pH 8, 0.3 M NaCl, 10 mM imidazole, 0.5 mg/mL lysosyme). After sonication, unbroken cells were removed by centrifugation at 12,000□x g for 15□min at 4□°C. The supernatant was loaded onto HisTrap Column (Cytiva) equilibrated with Buffer A (50 mM Tris-HCl pH 8, 0.3 M NaCl, 10 mM imidazole), and eluted with Buffer B (50 mM Tris-HCl pH 8, 0.3 M NaCl, 500 mM imidazole). Fractions of interest were then concentrated using a 30,000 Da molecular weight cut off (Amicon, Millipore) and loaded onto a Superdex 200 Increase 10/300 GL column (Cytiva) equilibrated with 50□mM Tris-HCl pH 8.0, 0.3 M NaCl.

Fractions containing ColE1-His6 were collected and concentrated as mentioned above. 7 mg of purified ColE1 was sent to GeneCust company to immunize New Zealand rabbits. Immunoglobulins were purified on an antigen affinity chromatography. Antibodies were kept at -20°C in PBS supplemented with 0,02% NaN3. Purified antibodies were tested on WT and Δ*colE1* mutant strains to assess their purity.

#### Anti-*S. sonnei* O-Antigen

*S. sonnei* CIP106347 was grown O/N in 5 mL LB medium at 37°C under shaking. The following day, *S. sonnei* was diluted to OD600 = 0.5 and grown to OD600 = 1.5. Bacteria were pelleted at 3200g 4°C for 15 minutes and incubated in 200 µL Tris 60 mM pH 8.0 supplemented with 200 µg/mL lysozyme (Carl Roth) for 30 minutes at 37°C. The suspension was supplemented with 200 µL 6% SDS suspended in Tris 60 mM pH 8.0 and boiled at 100°C for 10 minutes. Proteinase K was added at 50 µg/mL and incubated overnight at 37°C. After boiling at 100°C for 10 minutes, an additional 50 µg/mL Proteinase K were added and incubated at 55°C for 3 hours and boiled a final time at 100°C for 10 minutes. O-Antigen profiles were validated by SDS-PAGE followed by silver staining. Purified O-Antigen was sent to GeneCust company to immunize mice, using the same process as anti-ColE1. Purified antibodies were tested on *S. sonnei* CIP106347 smooth colony, as well as negative controls *S. sonnei* CIP106347 rough colony and *S. flexneri* 5a M90T.

### Ascorbate-deficient guinea pig model of shigellosis

Tissues from our previous study ^62^ were used for immunofluorescence stainings. After washing the slides in PBS and PBS + 0.1% Triton X-100, conjugated antibodies were directly added for 1 hour at room temperature in the dark, diluted in PBS + 0.5% Triton X-100 + 1% Bovine Serume Albumin (Euromedex). The following antibodies were used: Chembright Coralite488-conjugated (ProteinTech) 1 µg/mL mouse anti-O-Antigen *S. sonnei* (this study), 1 µg/mL Lightning Link AlexaFluor550-conjugated (Abcam) rabbit-anti-SigA (Debande et al., 2024), 1 µg/mL Lightning Link AlexaFluor647-conjugated (abcam), 1 µg/mL mouse anti-AmyloidBeta (clone WO2 ; ThermoFischer) and 1 µg/mL DAPI (Euromedex). Slides were washed again, mounted with ProLongGold® (Invitrogen), and imaged using a laser-scanning TCS SP8 confocal microscope (Leica).

### Biofilm formation

*Shigella* biofilms were grown as described by (Nickerson at al.^8^ with a few adjustments. Briefly, biofilms were grown from O/N cultures diluted 1:50 in LB + 0.25% Glucose supplemented or not with 0.4% bile salts (Sigma Aldrich). Glucose and bile salts powders were dissolved directly in LB the day of each experiment. After biofilm growth, OD600 was measured. Media were discarded and biofilms were washed with PBS then fixed with 4% PFA for 30 minutes. Biofilms were then stained with crystal violet and treated as described by Nickerson et al. ^63^. All experiments contained blank media, and all media were supplemented with streptomycin 50 µg/mL.

### Bactericidal killing assay

LB agar plates were overlaid with soft LB agar (0.7 % w/v) mixed with stationary phase indicator bacteria (*E. coli* BW25113, *E. coli* JW5503-1 Δ*tolC* and *P. megaterium*).

Bacteria strains that were tested for colicin activity were induced at OD600 = 1 using 0.5 µg/mL Mitomycin overnight, centrifuged at 9000 x g for 5min, 4° C, and the supernatant was sterile filtered (0.22 μm filters).

Filtered supernatants from the tester strains, purified ColE1, or purified F-SigA from different mutant *S. sonnei* strains (pSigA3, Δ*sigA* pSigA3, Δ*colE1* pSigA3 and Δ*sigA*) were then spotted in 2-μL drops onto the indicated strains’ lawns, the plates were incubated overnight at 37°C, and scanned to visualize the inhibition zone.

### Bioinformatic analysis of the distribution of SPATEs in a clinical dataset

*Shigella* genomes used in this study were previously described ^64–67^ . Sequence reads from this study were downloaded from the European Nucleotide Archive (Supplementary Table 4), adaptors and low-quality bases were trimmed with Trimmomatic v0.39 ^68^ . Sequence read quality was assessed with FastQC v0.11.6 (https://www.bioinformatics.babraham.ac.uk/projects/fastqc/) and MulitQC v1.8 ^69^ . Draft genome sequences were then assembled using Unicycler v0.4.9 ^70^ with - min_fasta_length set at 200 and annotated using Prokka v1.14.0 ^71^ . A pangenome of each *Shigella* spp. was constructed from the resulting GFF annotation files using Panaroo v1.5.1 ^72^, with default settings. The distribution of SPATE genes across the pangenome were assessed by first identifying the corresponding locus IDs for sigA (SF2968 & SSON_3223), sepA (SP_p0070), pic (SF2973), sat (SF2457T_3873) in the *S. flexneri* 2a strains 301 reference genome (accession: https://www.ncbi.nlm/nih.gov/nuccore/NC_004337) and 2457T (accession: AE014073.1), as well as the *S. sonnei* Ss046 reference genome (accession: CP000038.1), then matching these to homologous gene clusters in the pangenome to determine their presence, absence and distribution.

### SPATE structural analysis using AlphaFold and AmylPred2

50 dimeric models of the secreted SigA sequence were generated using AlphaFold-3. A distance matrix is subsequently computed based on pairwise structural alignment using TM-align. The distance between models is derived as the negative logarithm of the TM-score. Hierarchical clustering is performed on this matrix using the implementation provided in the SciPy library. This procedure is validated by visually inspected of the resulting clusters using PyMol. For each cluster, the medoid structure is extracted and designated as the representative dimer of its corresponding group. There representative dimers are then combined pairwise to model the formation of higher-order oligomers. Polymer growth is conducted by adding a new SigA monomer iteratively by structural alignment (TM-align) between one medoid dimer chain and the last chain of the growing polymer. After each addition, a clash score is computed to assess steric compatibility. Upon strict clash score increase, the growth process is halted, and the two dimerization interfaces are considered incompatible. Polymer assembly continued until a maximum of 20 monomers is reached. AmylPred2 predictions were conducted directly on the AmylPred2 website, and the data were visualized using Python.

### Mass–Spectrometry Analysis by NanoLC–MS/MS

For LC–MS/MS analysis of the in-gel samples, protein bands were excised from the SDS–PAGE gel and destained overnight at 4 °C in 50 mM ammonium bicarbonate containing 50% (v/v) acetonitrile. The gel pieces were subsequently dehydrated in 100% acetonitrile and incubated with 10 mM dithiothreitol for 45 min at 56 °C to reduce disulfide bonds. Cysteine residues were then alkylated with 55 mM iodoacetamide for 45 min at room temperature in the dark. After an additional dehydration step, the gel pieces were rehydrated with 50 ng of sequencing-grade modified trypsin (Promega) prepared in 50 mM ammonium bicarbonate and incubated overnight at 37 °C.

Peptides were sequentially extracted from the gel pieces using 60% acetonitrile containing 5% formic acid, followed by 100% acetonitrile. The peptide-containing supernatants were pooled and vacuum-dried using a SpeedVac concentrator. Dried peptides were resuspended in 0.1% formic acid and analyzed using a nanoElute 2 liquid chromatography system coupled to a timsTOF Pro 2 mass spectrometer (Bruker). Each sample was loaded onto a C18 trap column (Acclaim PepMap, 20 mm × 75 µm inner diameter, 3 µm particle size, nanoViper; Thermo Fisher Scientific) and separated on an Aurora Ultimate CSI C18 analytical column (25 cm × 75 µm inner diameter, 1.7 µm particle size; IonOpticks) using a 60-min gradient of solvent B, consisting of 0.1% formic acid in acetonitrile. The data were acquired in DIA-PASEF (Data-Independent Acquisition–Parallel Accumulation–Serial Fragmentation) mode of acquisition.

Mass spectrometry data were analyzed in library-free mode using DIA-NN software (version 2.3). Database searches were performed against the UniProt Shigella sonnei proteome database (4,068 sequences; release 2026_02) and the UniProtKB/Swiss-Prot Escherichia coli K-12 proteome database (4,403 sequences; release 2026_02). Precursor- and protein-level identifications were filtered at a false discovery rate of 1%. Match-between-runs and signal normalization were disabled. The deposition of the data from the in-gel samples will be submitted to the ProteomeXchange Consortium via the PRIDE partner repository; the validation process is ongoing.

### Statistical Analysis

The Shapiro-Wilk test was performed on samples with n>10. Samples that didn’t pass normality were analysed using the Mann-Whitney test. Samples with n<10 were analysed using Welch’s t-test. All statistical tests were performed using GraphPad Prism v11.0.0.

## Acknowledgments

A.S. and L.D. were granted with a PhD fellowship from the University of Strasbourg. We acknowledge support from the Biological Mass Spectrometry core facility at the University of Strabourg. This work was supported by the Agence Nationale de la Recherche ANR-22-CE15-0026 (DIFOX), ANR-22-CE15-0041-01 (NEUTROGAG) and ANR-25-CE15-4890 (BACTOCLOT). It has also benefitted from the support provided by the University for Advanced Study (USIAS) for a fellowship, within the French national program “Investment for the future” (IdEx-UNISTRA) (B.S.M.). This work was also supported by the Fondation pour la Recherche Médicale (FMR) (grant Equipe FRM #EQU202503020044).

## SUPPLEMENTARY INFORMATIONS

### Supplementary Figures

**Figure S1.**
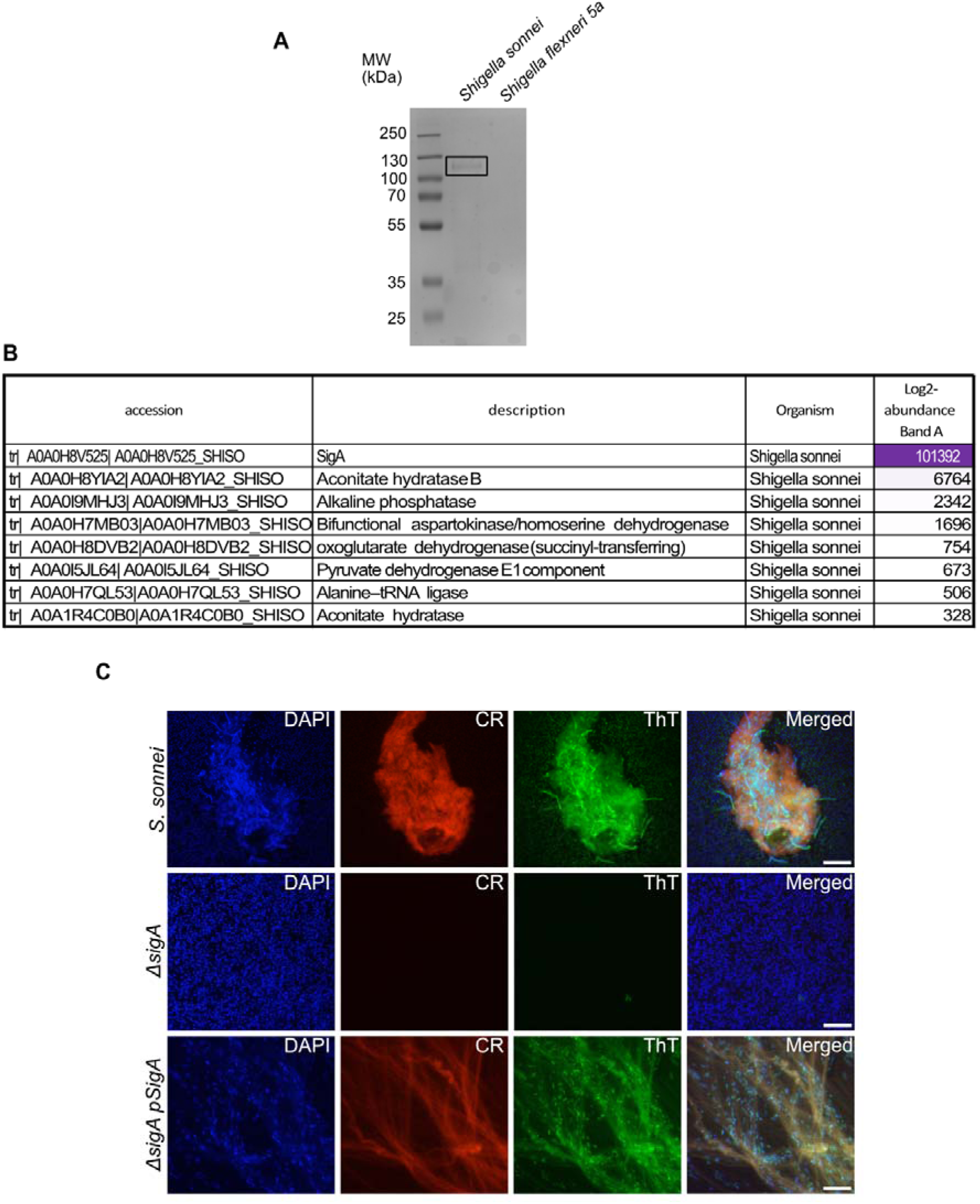
Identification of the protein involved in fibril formation as SigA. (**A**) SDS-PAGE gel separation of *S. sonnei* and *S. flexneri* 5a treated supernatant, corresponding to the sample shown in Fig. 1C. The squared band was analysed by Mass Spectrometry, as shown in (B). (**B**) Most abundant proteins in the sample shown in (A) were identified by mass spectrometry from the *Shigella sonnei* database. (**C**) Secretion of SigA amyloid fibrils by *S. sonnei* wild-type, D*sigA* and D*sigA* pSigA strains. DNA was stained with DAPI (blue), amyloid fibrils were stained with Congo red (CR, red) and ThioflavinT (ThT, green), and samples were analyzed by epifluorescence microscopy 40X. Scale bars, 10 mm.

**Figure S2.**
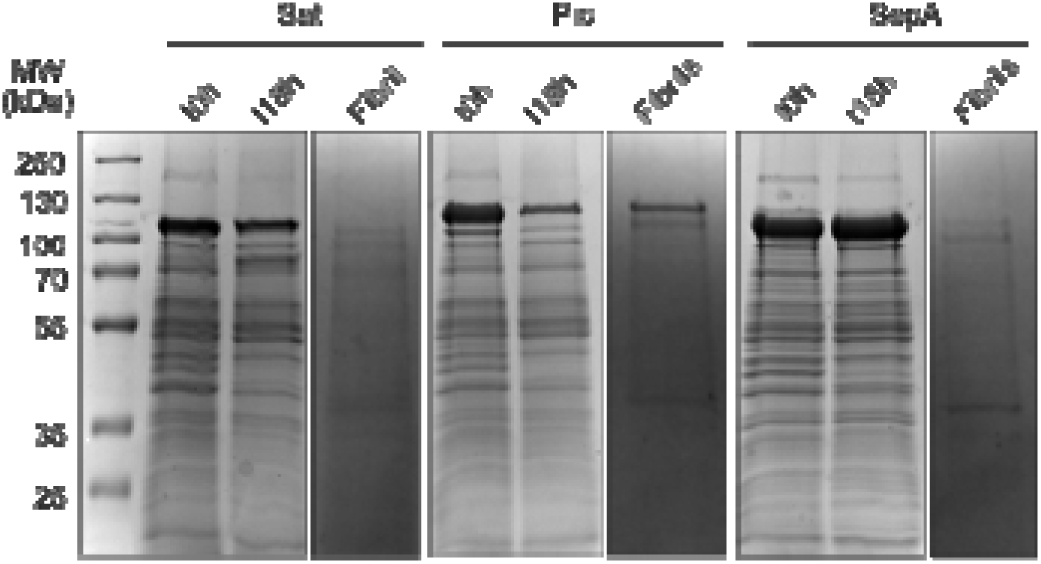
Pic is more prone to form amyloid fibrils. The protocol represented in Fig. 3A was applied to *E. coli* HB101 pSat, pPic and pSepA. As described in Fig. 3B, Purified fibrils, if produced, were resuspended in Urea 8M, separated on SDS-PAGE gel and stained with Coomassie.

**Figure S3.**
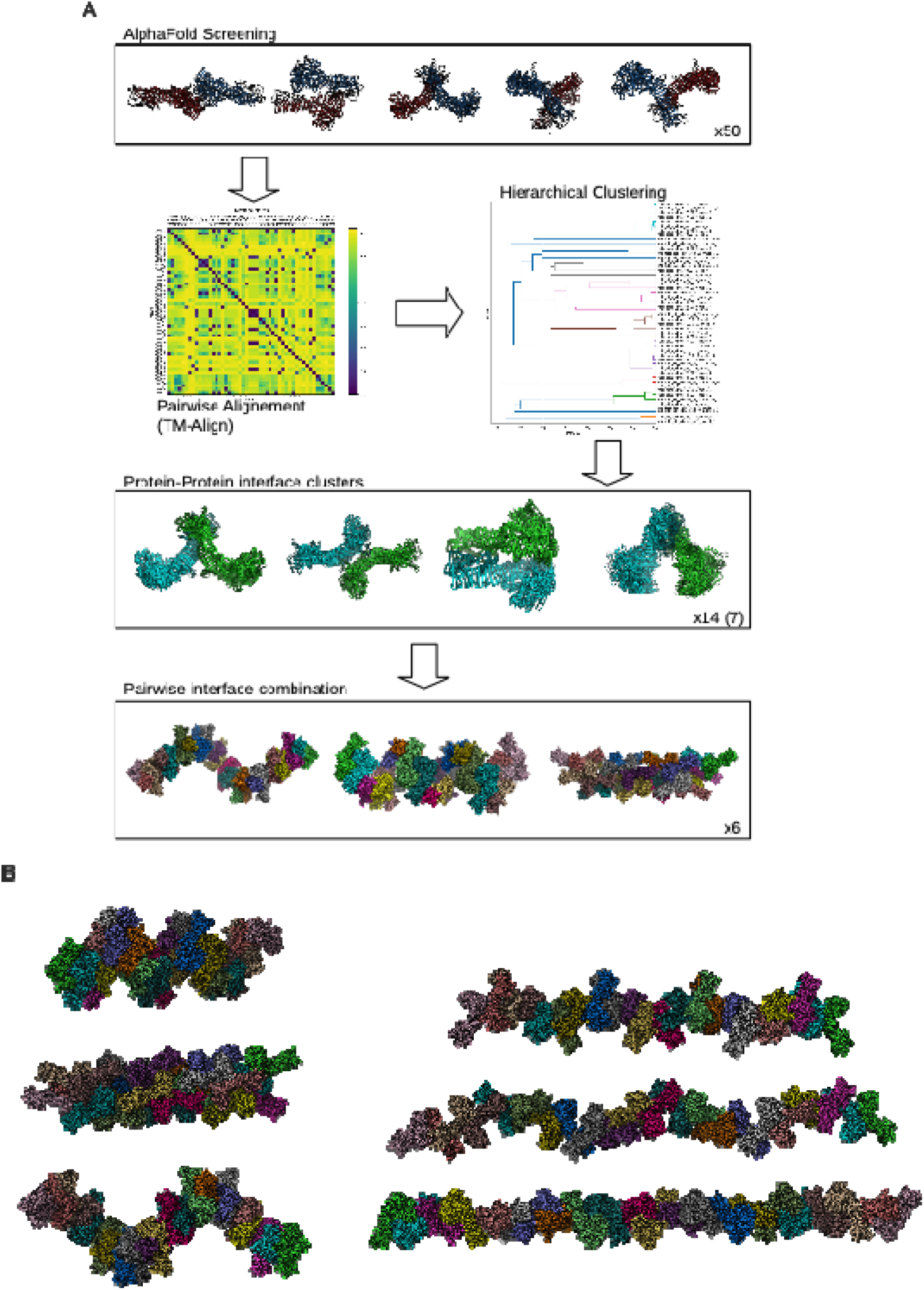
AlphaFold3 predicts that SigA forms rigid body filaments. (**A**) Description of the SigA rigid-body polymer modelling workflow 50 dimeric SigA structures were modelled using AlphaFold-3 and subsequently clustered. Following a pairwise comparison with TM-align and a hierarchical clustering procedure, using -log(TM-score) as distance. After cluster visual inspection (PyMol), the medoid structures are combined pairwise following an iterative structural alignment procedure (TM-align), resulting in growing polymers. (**B**) Six possible polymeric arrangements for SigA fibrillation, each shown with individual SigA monomers colored differently

**Figure S4.**
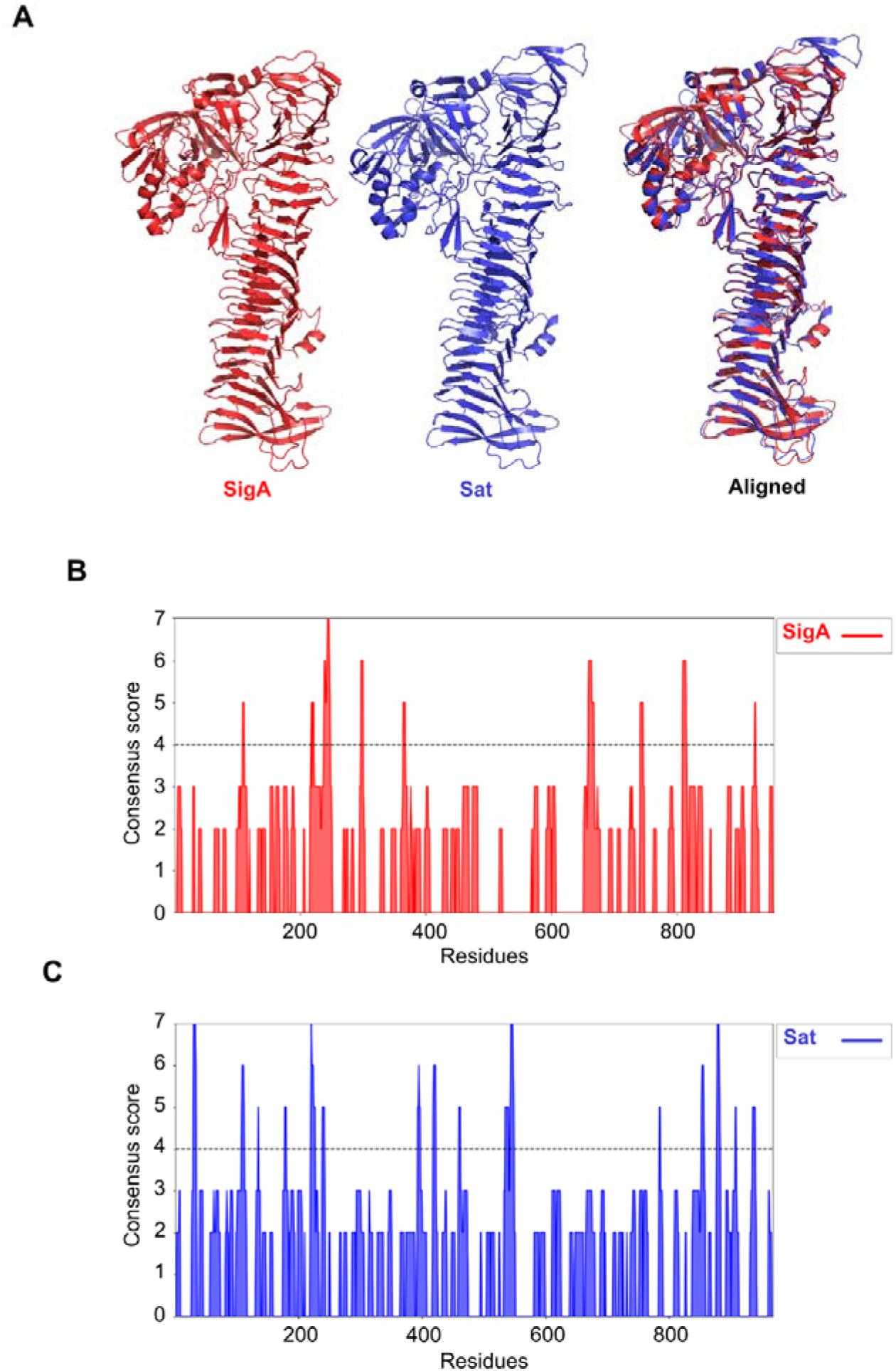
Comparison of SigA and Sat structures and predicted amyloid profiles. (**A**) SigA (ma-s7yqq) (red) and Sat (AF-Q8FDW4-F1-v6) (blue) passenger domains were aligned using PyMOL. (**B**) AmylPred2 amyloid hotspot prediction within the SigA passenger domain sequence (consensus score >4 is associated with a putative amyloid hotspot). (**C**) AmylPred2 amyloid hotspot prediction within the Sat passenger domain sequence (consensus score >4 is associated with a putative amyloid hotspot).

**Figure S5.**
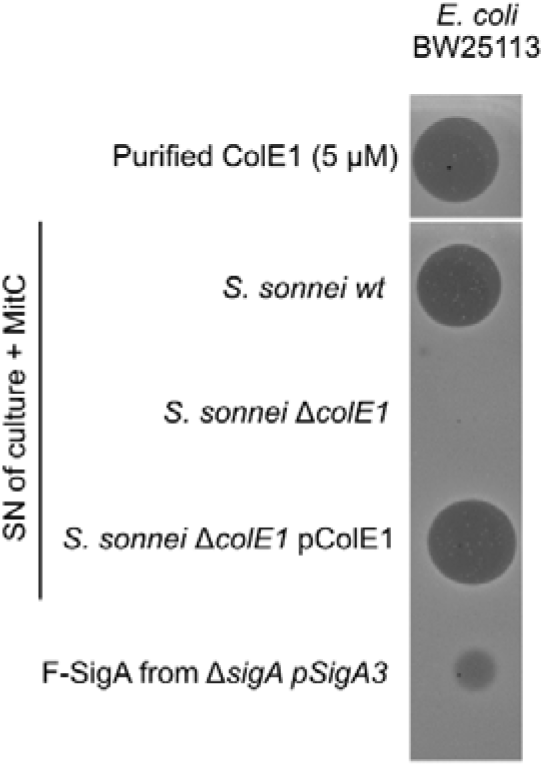
Purified F-SigA inhibits *E. coli* growth. *E. coli* BW25113 WT lawn killing assays on soft agar against purified ColE1 (5 μM), culture supernatant (SN) of *S. sonnei* WT, *ΔcolE1*, and *ΔcolE1* pColE1 strains treated with Mitomycin C (MitC, 0.5 µg/mL) or F-SigA from *S. sonnei* Δ*sigA* pSigA3.

**Table S1.** Bacterial strains.

| Strain | Description | Antibiotic Resistance/<u>Inductor</u> | Origin |
| --- | --- | --- | --- |
| *Shigella sonnei* | Clinical Isolate (CIP106347) | Streptomycin | Pasteur Institute Collection |
| *S. sonnei* $\Delta sigA$ | *sigA* mutant | Streptomycin/<br>Chloramphenicol | Debande *et al.*, 2024 |
| *S. sonnei* $\Delta sigA$ pSigA | *sigA* mutant complemented with pAS1-*sigA* | Streptomycin/<br>Chloramphenicol/<br>Ampicillin | This study |
| *S. sonnei* $\Delta sigA$ pSigA<sub>S258A</sub> | SigA(S258A) expression under *sigA* native promoter | Streptomycin/<br>Chloramphenicol/<br>Ampicillin | This study |
| *S. flexneri* 5a M90T-Sm | WT Strain | Streptomycin | Lab collection |
| *S. flexneri* 2a 2457T | WT Strain | None | Lab collection |
| *Escherichia coli* CFT073 | UPEC WT strain | None | From Eric Oswald |
| *E. coli* HB101 | Lab strain |  | Lab collection |
| *E. coli* HB101 pSigA3 | Recombinant SigA overexpression through pUC19 | Ampicillin<br><u>IPTG</u> | This study |
| *E. coli* HB101 pSigA | SigA expression under *sigA* native promoter | Ampicillin | This study |
| *E. coli* HB101 pSigA<sub>S258A</sub> | SigA(S258A) expression under *sigA* native promoter | Ampicillin | This study |
| *E. coli* HB101 pSat | Sat expression under *sigA* native promoter | Ampicillin | This study |
| *E. coli* HB101 pPic | Pic expression under *sigA* native promoter | Ampicillin | This study |
| *E. coli* HB101 pSepA | SepA expression under *sigA* native promoter | Ampicillin | This study |
| *S. sonnei* $\Delta sigA$ pSigA3 | Complemented mutant expressing SigA under Lac promoter | Streptomycin /<br>Chloramphenicol/<br>Ampicillin / <u>IPTG</u> | Lab collection |
| *E. coli* BW25113 | WT strain | None | from Nicholas Housden : KEIO collection |
| *E. coli* JW5503-1 | F-, $\Delta(araD-araB)567$, $\Delta lacZ4787(::rrnB-3)$, λ-, $\Delta tolC732::kan$, rph-1, $\Delta(rhaD-rhaB)568$, hsdR514 | Kanamycin | from Nicholas Housden : KEIO collection |
| *Priestia megaterium* | WT strain | None | Kind gift from Dr. Isabelle Caldelari (CNRS) |
| *S. sonnei* $\Delta colE1$ | *colE1* mutant | Streptomycin / Chloramphenicol | This study |
| *S. sonnei* $\Delta colE1$ pColE1 | *colE1* mutant overexpressing *colE1*, *immunity* and *lysis* genes under *colE1* promoter | Streptomycin / Chloramphenicol/ Ampicillin | This study |
| *S. sonnei* $\Delta colE1$ pSigA3 | *colE1* mutant overexpressing SigA under Lac promoter | Streptomycin / Chloramphenicol/ Ampicillin / <u>IPTG</u> | This study |
| *E. coli* Nico21 pBR03 | Lab strain expressing ColE1-His6 under Lac promoter | Kanamycin / <u>IPTG</u> | This study |

**Table S2.** Plasmids.

| Plasmid | Description | Antibiotic Resistance/<u>Inductor</u> | Origin |
| --- | --- | --- | --- |
| pSigA | pBBR1-SigA. pBBR1-MCS4 backbone (Low copy), *sigA* promoter, gene and terminator from *S. sonnei* CIP106347 | Ampicillin | This study |
| pSigA<sub>S258A</sub> | pBBR1-*sigA*<sub>S258A</sub>. pBBR1-*sigA* with a point mutation transforming the Ser258 into Ala | Ampicillin | This study |
| pSat | pBBR1-*sat*. *sigA* promoter and *sat* gene from UPEC CFT073 | Ampicillin | This study |
| pPic | pBBR1-*pic*. *sigA* promoter, *pic* gene modified from *S. flexneri* 2a *pic* gene | Ampicillin | This study |
| pSepA | pBBR1-*sepA*. *sigA* promoter and *sepA* gene from *S. flexneri* 5a M90T-Sm | Ampicillin | This study |
| pSigA3 | pUC19 backbone (high copy) Lac promoter and *sigA* gene from *S. flexneri* 2a 2457T | Ampicillin<br><u>IPTG</u> | Meza-Segura et al., 2021 |
| pBR03 | pET28a-*colE1*-His6X (C-ter) | Kanamycin<br><u>IPTG</u> | This study |
| pColE1 | pUC19; *colE1* promoter and *colE1*, *immunity* and *lysis* genes from *S. sonnei* | Ampicillin |  |
| pTwist-*pic* | pUC19 vector expressing a synthetic *pic* fragment with alternative codons that do not co-encode a *set1AB* toxin | Ampicillin | This study |

**Table S3.**
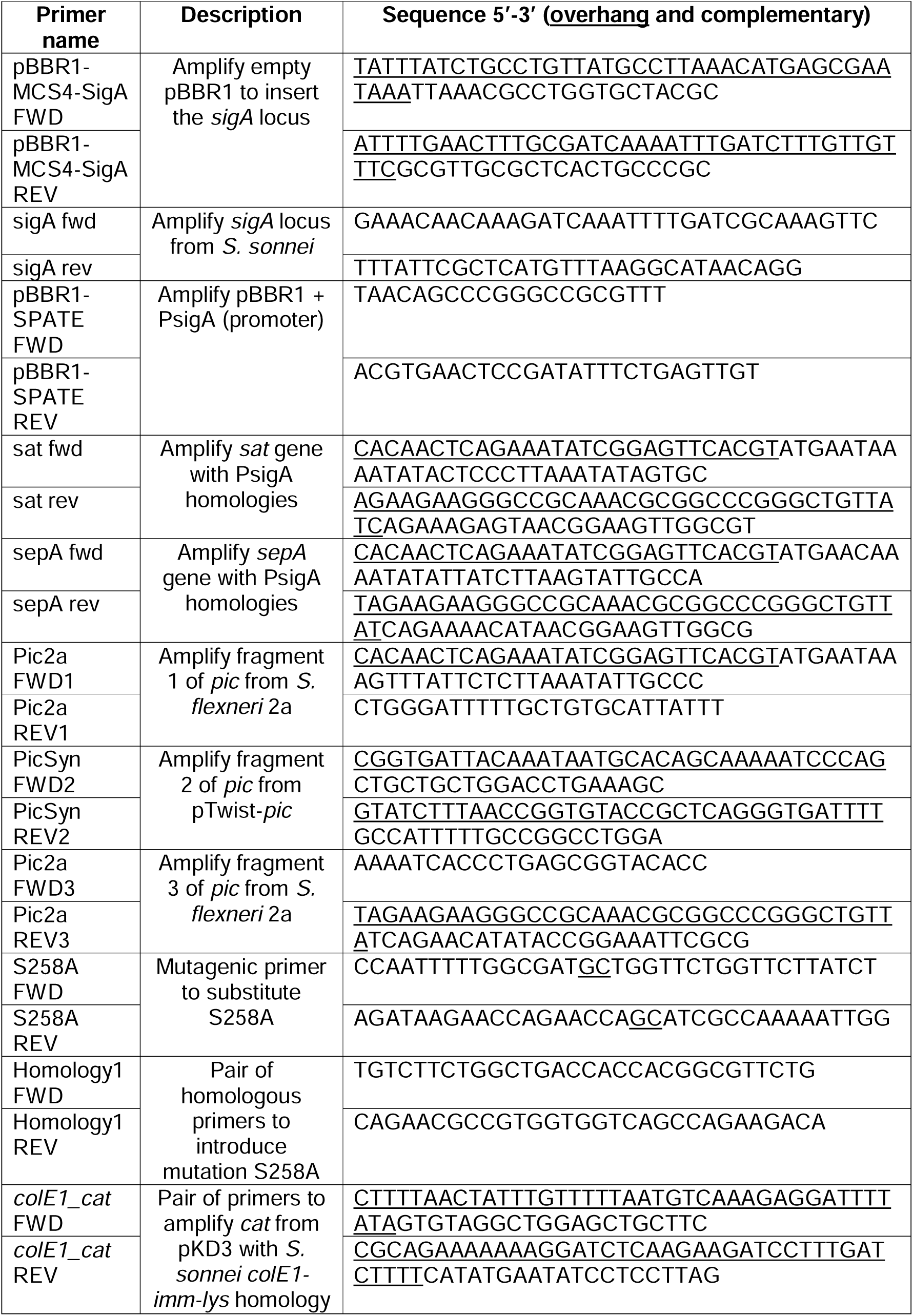
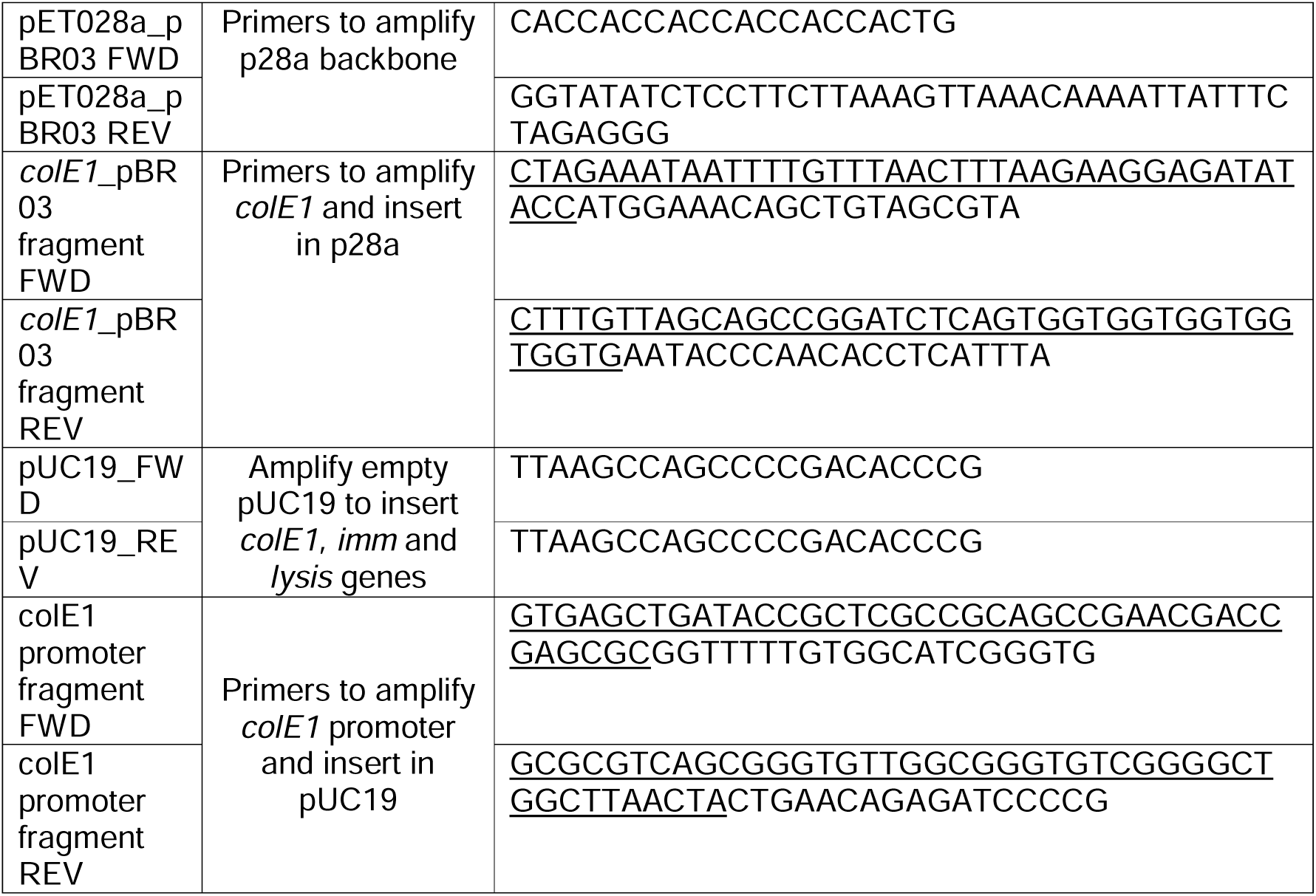
Primers.

